# A novel lineage of polyomaviruses identified in bark scorpions

**DOI:** 10.1101/2021.05.12.443864

**Authors:** Kara Schmidlin, Simona Kraberger, Chelsea Cook, Dale F. DeNardo, Rafaela S. Fontenele, Koenraad Van Doorslaer, Darren P. Martin, Christopher B. Buck, Arvind Varsani

## Abstract

Polyomaviruses are non–enveloped viruses with circular double-stranded DNA genomes that range in size from ∼4–7 kilobasepairs. Initially identified in mammals, polyomaviruses have now been identified in birds and a few fish species. Although fragmentary polyomavirus-like sequences have been detected as apparent ‘hitchhikers’ in shotgun genomics datasets for various arthropods, the possible diversity of these viruses in invertebrates remains unclear. In general, polyomaviruses are host-specific, showing strong evidence of host-virus co–evolution. Identification of polyomaviruses in a broader range of animals could shed useful light on the evolutionary history of this medically important group of viruses. Scorpions are predatory arachnids that are among the oldest terrestrial animals. Thus far, viromes of arachnids have been under–sampled and understudied. Here, high–throughput sequencing and traditional molecular techniques were used to explore the diversity of circular DNA viruses associated with bark scorpions (*Centruroides sculpturatus*) from the greater Phoenix area, Arizona, USA. The complete genomes of eight novel polyomaviruses were identified. Analysis of *Centruroides* transcriptomic datasets elucidated the splicing of the viral late gene array, which is more complex than that of vertebrate polyomaviruses. Phylogenetic analysis provides further evidence of co-divergence of polyomaviruses with their hosts, suggesting that at least one ancestral species of polyomaviruses was circulating amongst the primitive common ancestors of arthropods and chordates.

## Introduction

Scorpions are one the oldest terrestrial arthropods, with fossil records dating back 450 million years. They are venomous, predatory animals that belong to the class Arachnida. Nearly 2200 scorpion species are recognized, occurring in a wide range of ecosystems including deserts, grasslands, savannahs, deciduous forests, pine forests, intertidal zones, rain forests and caves (1). Their bodies are separated into two segments, a cephalothorax and an abdomen that includes the tail and stinger, four pairs of legs and pedipalps with plier–like pinchers (2). Bark scorpions (*Centruroides sculpturatus*) are nocturnal, generalist predators that consume a wide variety of insects, spiders, centipedes and other scorpions. They inhabit arid desert and riparian regions in south–western North America (2, 3). Although these scorpions are desert adapted, water is a limiting factor, so they are most commonly found in riparian areas. The bark scorpion is a common inhabitant of modern desert cities (4).

The first polyomavirus was discovered in the mid-1950s as a filterable agent and was shown to induce multiple classes of tumors when experimentally injected into animals, suggesting the name poly (multiple) oma (tumors). Polyomaviruses have been recovered from a variety of mammals including humans, bats, and rodents, and diverse polyomavirus species have recently been reported in an assortment of birds and fish (5, 6).

Polyomaviruses have a conserved genome organization that consists of an early region and a late region separated by a regulatory region (RR) that encompasses the early and late promoters as well as the origin of replication. Polyomavirus genomes are remarkably simple, encoding only three absolutely essential proteins. The Large Tumor antigen (LT) is a multipurpose protein that encodes a superfamily 3 helicase domain that unwinds the origin of replication and primes the host cell DNA replication machinery to replicate the viral genome. LT also encodes a DNAJ-like domain, which encodes a hallmark HPDKGG motif, and a pRB-interaction motif, LXCXE (7). All known polyomavirus late regions encode a major capsid protein, VP1, which forms the non-enveloped icosahedral surface of the virion, and a minor capsid protein, VP2, that associates with the inner surface of the assembled virion. A hallmark feature of VP2 is an N-terminal myristoylation motif. (7).

Although polyomaviruses are host specific, strict virus-host co-divergence models do not fully explain polyomavirus evolution. A growing body of evidence suggests an ancient and relatively stable association of polyomaviruses with their hosts (5). While this long–standing association implicates co–divergence as the main driver of polyomavirus evolution, there are deviations that indicate additional factors may occasionally contribute to their evolutionary history including switching between closely related animal hosts, intra-host lineage duplication followed by divergence (i.e., viral speciation within the same host species), and incomplete lineage sorting (persistence of viral polymorphisms during successive host divergence events) (5). Viral metagenomics and molecular techniques, such as degenerate PCR and random-primed rolling-circle amplification (RCA), are enabling the identification and recovery of polyomaviruses from novel hosts at an increased frequency. This has allowed for a deeper understanding of polyomavirus evolutionary history. Shotgun deep sequencing has also begun revealing viral sequences as inadvertent contaminants in genomics and transcriptomics studies. Since shotgun genomics studies are not designed to enrich for viruses and do not require careful management of sample cross-contamination, these adventitious virus sequences can be difficult to interpret. For example, although a prior data mining effort detected highly divergent polyomavirus-like sequences in a shotgun genomics dataset for California bark scorpion, it is difficult to rule out the possibility that the sequence came from an ectoparasite or environmental source, as opposed to a productive infection of the scorpion itself (5). Because the ultimate sources of data-mined sequences are uncertain, directed discovery of additional polyomavirus sequences in invertebrates could help further resolve the evolutionary history of this group. This information will help generate a taxonomic framework that can be utilized to accurately categorize novel viruses in such a way that sequence similarities could be used by researchers and clinicians to aid in predicting hosts, tissue tropism and disease states.

The International committee for the taxonomy of viruses (ICTV) recommends using phylogeny– based taxonomy of LT protein sequences to classify polyomaviruses into four recognized genera as well as several unassigned species (8, 9). The four established genera are *Alphapolyomavirus, Betapolyomavirus* and *Deltapolyomavirus* which contain species that infect mammalian hosts and *Gammapolyomavirus* which contains species that infect avian hosts. Recently identified polyomaviruses associated with fish datasets (5, 6, 10) have not yet been assigned to a genus. In the present report we identify and highlight the diversity of eight complete polyomavirus genomes recovered from the viscera of carefully cleaned bark scorpion carcasses and propose a taxonomic assignment for the group.

## Material and methods

### Sample collection and preparation

Four bark scorpions were collected from two locations, Phoenix (n=3) and Tempe (n=1), Arizona, USA, between August and November 2018 and stored at –80°C. Scorpions were surface sterilized by washing in 95% ethanol for 15 seconds followed by a wash in 10% bleach for 60 seconds before rinsing three times in pure water. The specimens were allowed to air dry in a petri dish lined with sterile filter paper in a laminar flow hood prior to dissection. The stinger was removed and a sterile scalpel used to cut along the side of the body in between the legs and body plate. Sterile forceps were used to remove the top plate so the internal cavity was visible. The gut and liver were removed and each was placed in a sterile 1.5ml tube and homogenized in 500 µl of SM buffer (50 mM Tris–HCl, 10 mM MgSO_4_, 0.1 M NaCl, pH 7.5). The homogenate was processed as previously described in (11). Viral DNA was extracted using the Roche high pure viral nucleic acid kit (Roche, USA) according to manufacturer’s instructions and circular DNA was enriched by rolling circle amplification (RCA) with TempliPhi™ amplification kit (GE Healthcare, USA).

### High throughput sequencing, de novo assembly and recovery of viral genomes

The RCA products were used to prepare Illumina sequencing libraries which were sequenced on an Illumina HiSeq4000 sequencer (2×100 bp paired end library) and the reads were *de novo* assembled using SPAdes 3.12 (12). Contigs of >1000 nts were analyzed using BLASTx (13) against a local viral protein sequence database. Eight *de novo* assembled contigs were found to have sequence similarities to dsDNA viruses from the family *Polyomaviridae*.

Based on the contigs, three sets of abutting primers were designed (Table 1) to recover the complete circular viral genomes by PCR. For each PCR amplification, 1 µl of RCA product was used as a template with specific primer pairs and Kapa Hifi Hotstart Ready Mix (Kapa Biosystems, USA) using the following thermal cycling protocol: initial denaturation at 95°C for 3 min followed by 25 cycles 98°C for 20 s, 60°C for 15 s, 72°C for 6 min, final elongation at 72°C for 6 min and a final renaturation at 4°C for 10 min. The resulting ∼5 kb amplicons were resolved on a 0.7% agarose gel, excised, gel purified, ligated into pJET1.2 vector (ThermoFisher Scientific, USA) and transformed into XL1 Blue *Escherichia coli* competent cells. The resulting recombinant plasmids were Sanger sequenced at Macrogen Inc. (South Korea) by primer walking. The Sanger sequence contigs were assembled using Geneious Prime 2019.1.1 (Biomatter Ltd, New Zealand).

**Table 1:**
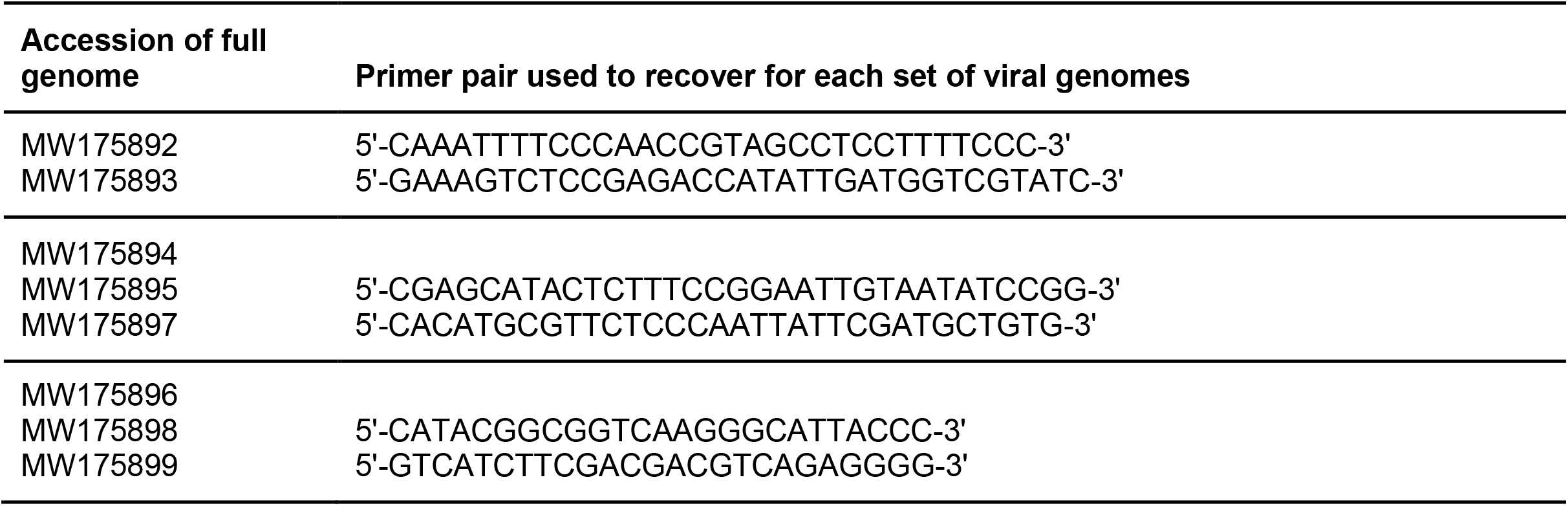
Primer details for each recovered scorpion polyomavirus

### Phylogenetic analysis

A curated dataset that includes representative sequences of all currently known polyomavirus species was downloaded on 10^th^ December, 2020 from https://ccrod.cancer.gov/confluence/display/LCOTF/Polyomavirus and used in all proceeding analyses. A maximum–likelihood of the aligned genomes of the polyomaviruses identified in this study together with two assembled polyomaviruses (LN846618 LN846619) from SRA SRS561933 using GTR+G substitution matrix. A maximum–likelihood phylogenetic tree was constructed using the LT amino acid sequences from eight scorpion-derived polyomavirus genomes from this study as well as those from LN846618-LN846619 together with those from the curated dataset and also those available in the public database GenBank as part of PRJNA642917 (SAMN13619660) and PRJDB7092 (SAMD00127334) from African social spider and Orb-weaving spider, respectively. We also included the LT-like sequence of circular molecule MH545547 (14) and that of bandicoot papillomatosis carcinomatosis viruses 1 and 2 (EU069819 and KP768176). The dataset was aligned using MAFFT (15) and a ML phylogenetic tree inferred using PHYML (16) with the model LG+G+I as determined by ProtTest (17). Branches with <0.8 aLRT branch support were collapsed using TreeGraph 2 (18) and the phylogenetic trees were visualized using IToL (19).

### Pairwise identities

Whole genome nucleotide sequences as well as LT and VP1 amino acid pairwise identities were determined using sequence demarcation tool (SDT) version 1.2 (20).

### Recombination analysis

The genomes of the arachnid polyomaviruses were aligned using MAFFT and this alignment was used to detect evidence of recombination using RDP5 (21) with default settings. Detected recombination signals with an associated p-value <0.05 for three or more of the seven recombination detection methods implemented in RDP5 were accepted as plausible evidence of recombination.

## Results and Discussion

### Genome organization and gene content

Four bark scorpions were collected in Phoenix (n=3) and Tempe (n=1), Arizona, dissected and analyzed for the presence of polyomaviruses. From three of the samples, eight complete genomes generally consistent with the organization of known polyomaviruses were recovered (Figure 1). Classic mammalian and avian polyomaviruses range from 4.7 to 5.4 kb in length. The scorpion polyomavirus genomes, in contrast, range from 5420 to 5945 bp. HHPred (22, 23) analyses and scans for hallmark linear motifs revealed open reading frames (ORFs) encoding homologs of LT, VP1 and VP2. The LT ORF, which lacks an ATG initiation codon, encodes an N-terminal DNAJ-like domain, an LXCXE motif, an endonuclease-like origin-binding domain, and a superfamily 3 helicase domain. Many polyomaviruses encode an alternative LT ORF (ALTO) protein overprinted in the LT +1 frame (24). A majority of the scorpion polyomaviruses encode an ALTO-like ORF. Well-studied mammalian polyomaviruses encode an Agno protein just upstream of VP2. In avian polyomaviruses the protein is incorporated into virions and is known as VP4 (25). Each of the scorpion polyomavirus genomes encodes a syntenic candidate Agno ORF. The VP2 ORF encodes a predicted N-terminal myristoylation signal (26) and HHPRED analyses suggest predicted C-terminal structural similarity to bacteriophage holin proteins. The central portion of the inferred VP1 ORF shows high-confidence hits for known polyomavirus VP1 structures.

**Figure 1:**
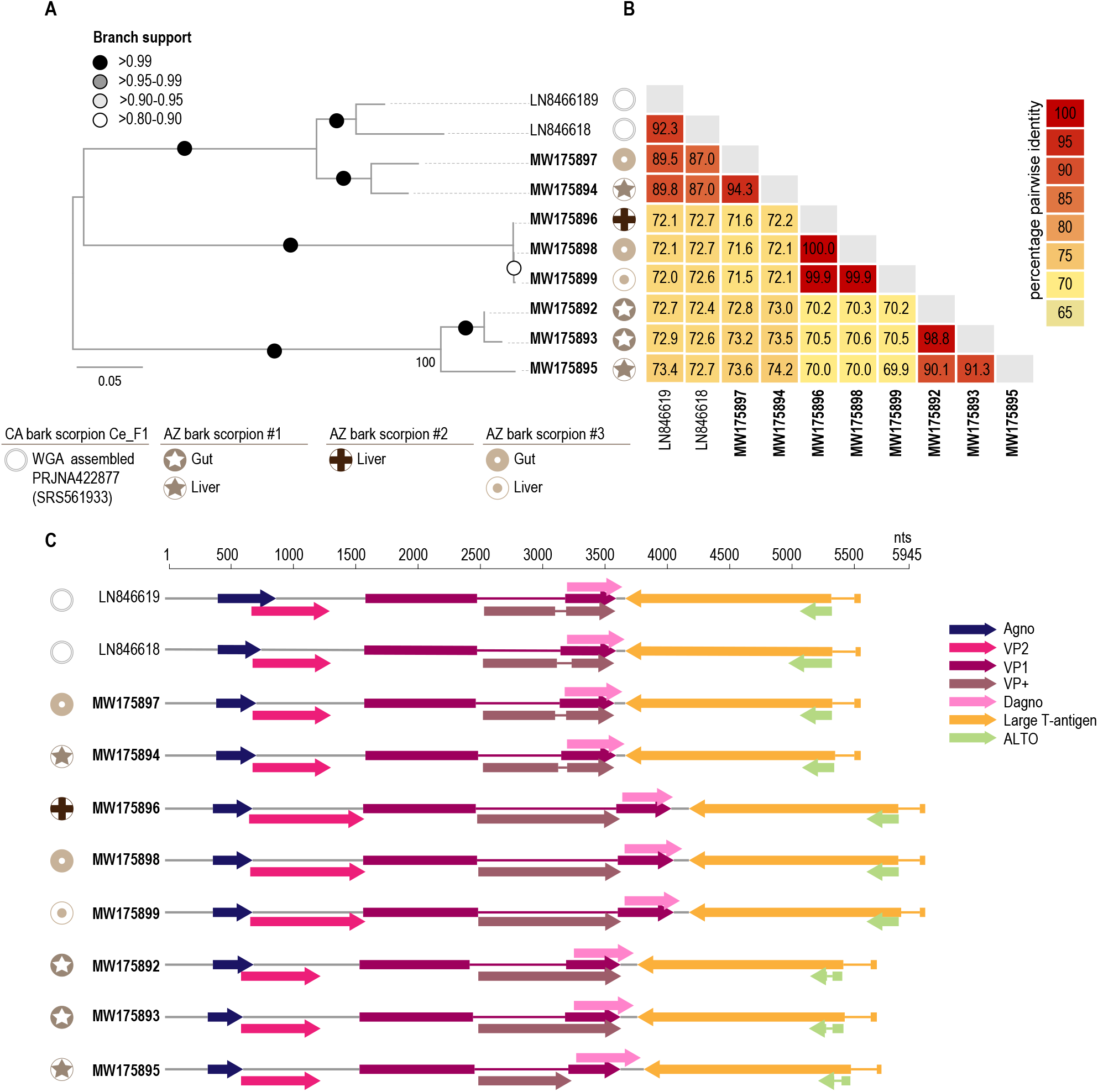
A. Complete genome maximum likelihood (ML) phylogenetic tree of nucleotide sequences identified in this study compared with the original WGS-assembled bark scorpion polyomavirus genome (5). Tissue sources of the recovered viral genomes are denoted by symbols. B. Genome-wide percentage pairwise identity comparisons between the scorpion polyomaviruses. C. Genome organization of the scorpion polyomaviruses with color coded-open reading frames.

The late regions of the scorpion polyomaviruses also show several unrecognizable ORFs. To better understand the expression of these proteins we performed data mining on available transcriptomics datasets for scorpions. A complete polyomavirus genome was assembled from RNAseq dataset SRR5958578 and was used to map introns. It was also possible to map a set of bark scorpion transcripts from BioProject PRJNA422877 to the previously data-mined WGS-assembled sequence LN846618. Although neither of the transcriptomic datasets had deep coverage of the early region, clear introns were identified in late region transcripts. The mapping revealed spliced transcripts encoding four possible isoforms of Agno, a C-extended VP1, and a protein with a predicted coiled coil structure that we name VP+. An additional ORF, which we name “downstream Agno” (Dagno), is overprinted in the +1 frame of VP+ (Figure 1). It seems conceivable that the more elaborate set of late proteins observed in the scorpion polyomaviruses might enable construction of a slightly larger virion, consistent with the larger genomes of these viruses.

### Similarity comparison

In bark scorpion carcass #1, four polyomaviruses were identified from the gut (n=2) and liver (n=2). The two from the gut share 99% genome-wide identity and the ones from the liver share 74% identity. Overall the four polyomavirus genomes share 73-99% identity (Figure 1). These AZ bark scorpion polyomaviruses share 73-90% genome-wide identity with the Baja California bark scorpion polyomavirus 1 (LN846619) originally assembled from WGS data (5).

In bark scorpion carcass #3, a total of three genomes were recovered from the gut (n=2) and liver (n=1). These share 72-100% genome-wide identity (Figure 1) in which 100% identical isolates were recovered from both tissue types. In AZ bark scorpion #2 only one genome was identified and it is 100% identical to the one identified in bark scorpion carcass #3 (from the liver and gut).

Official polyomavirus taxonomy guidelines (9) specify that polyomaviruses that share <85% LT pairwise nucleotide identity should be considered to belong to distinct viral species. By this standard, genomes recovered from bark scorpions can be assigned to three unique species (Figure 2). Polyomavirus sequences recovered from the gut and liver of bark scorpion samples #1 and #3, along with the original bark scorpion LN846619, represent species 1. Polyomavirus sequences that represent species 2 were recovered from bark scorpion samples #2 and #3, from both the liver and gut. Polyomavirus species 3 is comprised of sequences from bark scorpion #1 with isolates from both liver and gut tissues (Figure 2). The fact that identical or nearly identical viruses were recovered from necropsied tissue of multiple scorpions indicates that these viruses are endemic to these animals.

**Figure 2:**
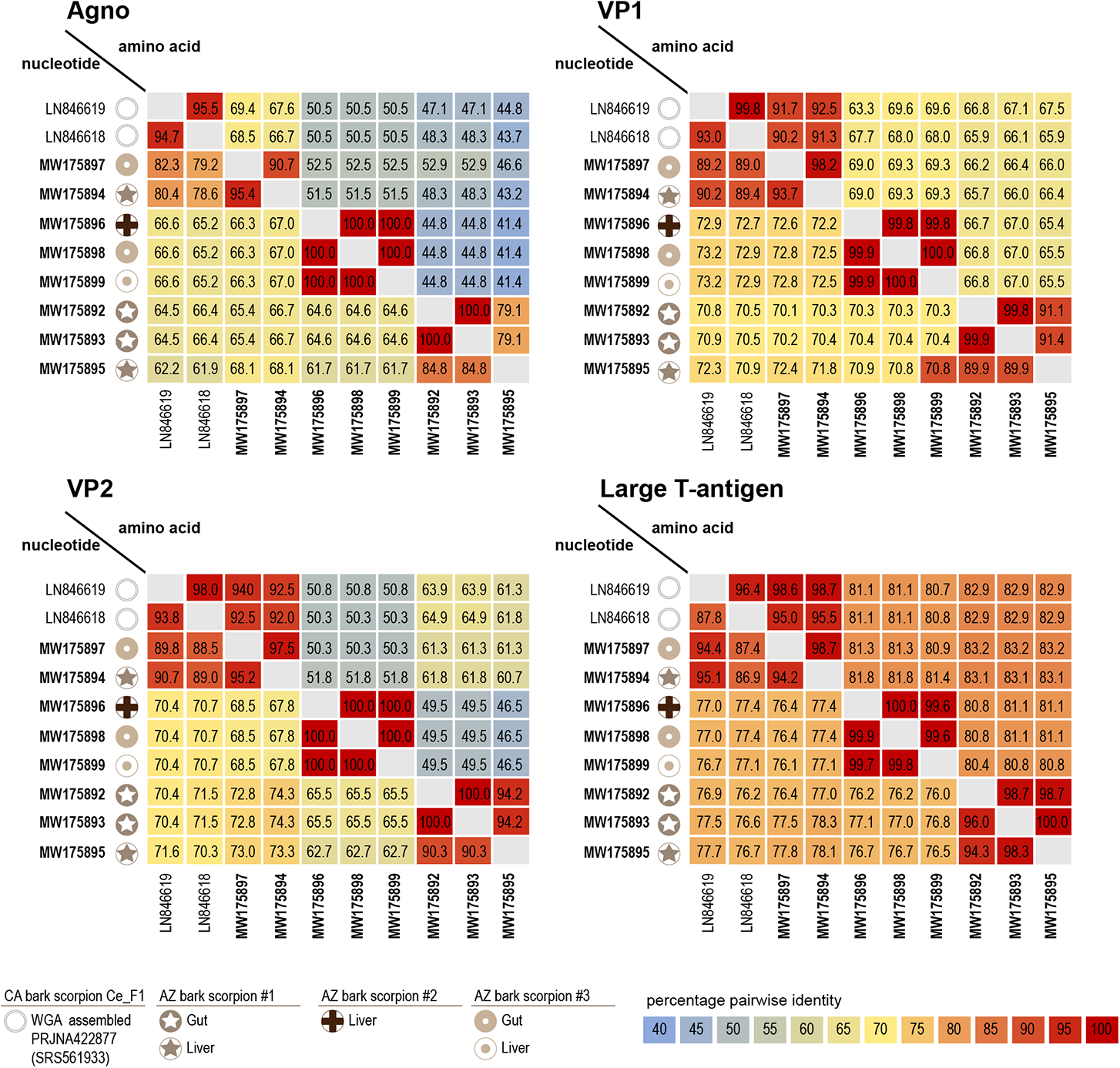
Percentage pairwise identities of four major polyomavirus ORFs. Nucleotide and amino acid comparisons are shown. The sources of the recovered viral genomes are denoted by symbols.

At the nucleotide level, the scorpion polyomaviruses share between 55 and 61% whole genome pairwise identity with all non-arachnid polyomaviruses. LT protein sequences of the scorpion polyomaviruses share between 21 and 35% amino acid pairwise identity with all other polyomaviruses and >80% identity amongst themselves. VP1 protein sequences show even higher diversity sharing only 19–31% identity with all other polyomavirus, >56% identity with each other (Figure 2).

### Phylogenetic analysis of LT

A maximum likelihood phylogenetic tree was constructed from LT amino acid sequences (5). Sequences included in this analysis were available prior to December 2020 and included putative LT sequences from a guitarfish polyomavirus, as well as LT-bearing polyomavirus/papillomavirus chimeric viruses found in bandicoots (27, 28). The phylogenetic tree (Figure 3) shows distinct clades that are generally consistent with expected phylogenetic placement of their host species within broad animal phylogeny. It was possible to completely collapse the polyomavirus clades derived from placental mammals and avians since these were monophyletic (with the exception of two divergent bat-associated polyomavirus genomes) (Figure 3). The clades containing fish-derived polyomaviruses branched basal to the mammalian and avian polyomavirus clades and roughly reflect the genetic diversity and evolutionary relationships of the fish hosts. Importantly, the scorpion derived sequences cluster together and branch from a location within the polyomavirus tree that might conceivably be the root of the tree. Furthermore, LT-like sequences have been identified in African social spider, giant hose spider, and orb-weaver spider that cluster with the scorpion polyomavirus LTs, suggesting a broad arachnid-specific clade (Figure 3).

**Figure 3:**
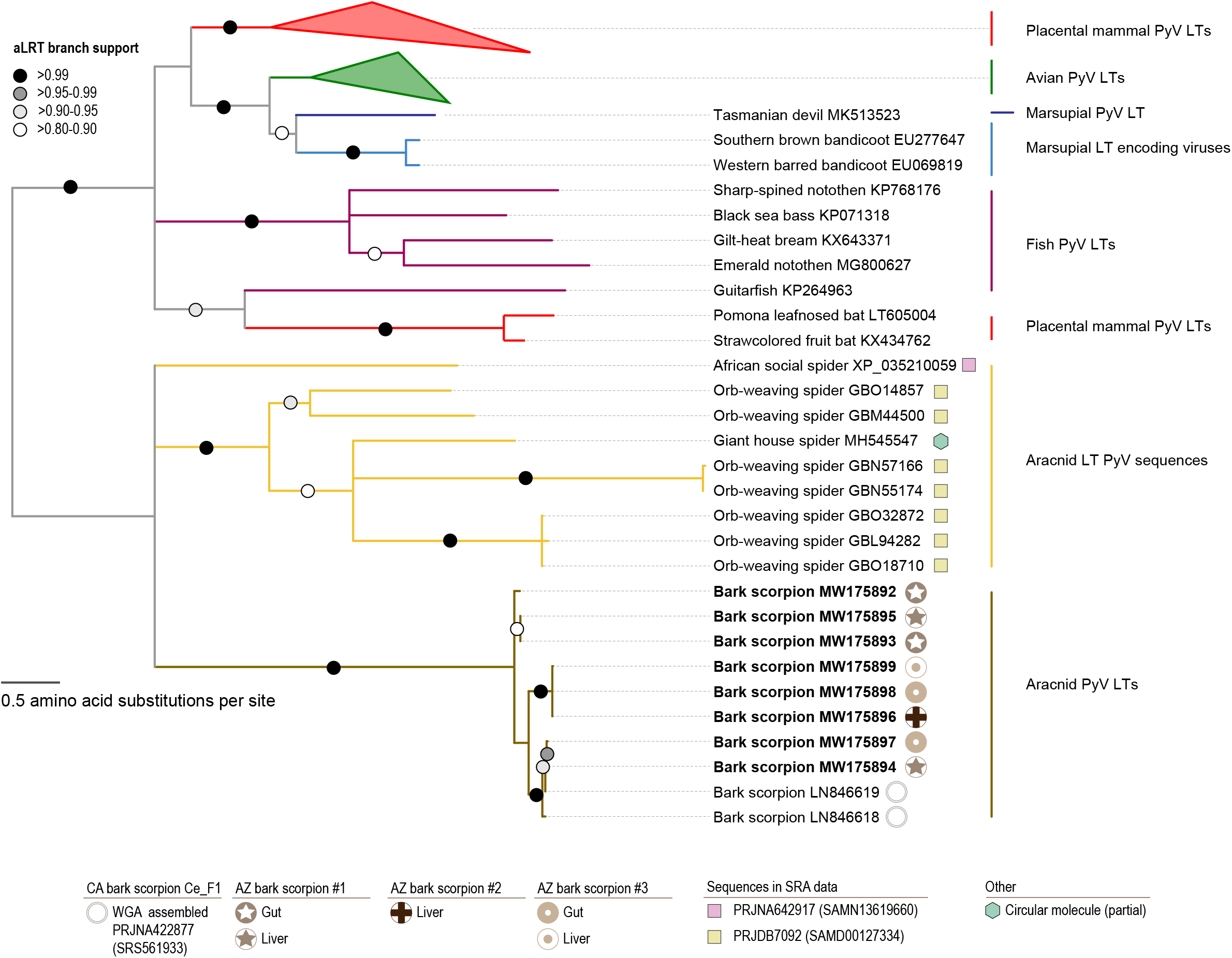
Mid-point rooted maximum likelihood phylogenetic tree of the LT sequences of known polyomaviruses, bandicoot papillomatosis carcinomatosis virus and those identified in this study. The core fish polyomavirus LTs are highlighted with purple clades, the unusual bat derived polyomavirus LTs are highlighted with orange clade. In the case of the scorpion polyomavirus LTs, the symbols represent the source.

### Recombination analysis

Two potential recombination events were detected in two of the scorpion polyomavirus sequences (summarized in Figure 4). Both the recombinant regions were identified in the LT. In the case of MW175893 (from the gut of the AZ bark scorpion #1), a recombinant region spanning 1031 nts appears to have been derived from a parental virus most closely resembling MW175895; a virus identified in the liver of the AZ bark scorpion #1. Whereas in the case of MW175897 (from the gut of the AZ bark scorpion #1) a much smaller, 64 nt long, recombinant region appears to have been derived from a virus most closely resembling MW175892, MW175893 and MW175895 (all of which were identified from AZ bark scorpion #1).

**Figure 4:**
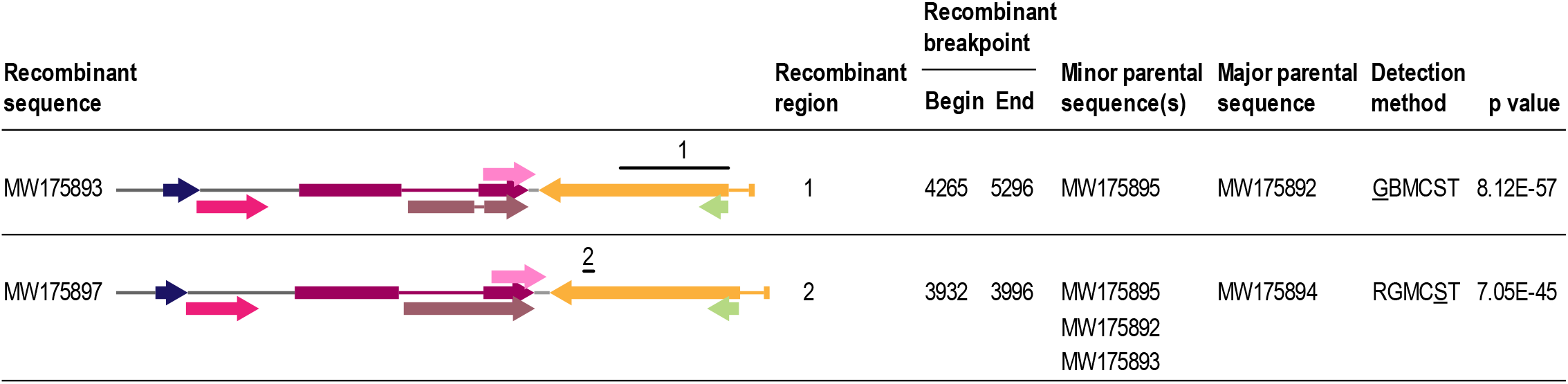
Summary of the recombination analysis of the scorpion polyomaviruses. The two recombination events detected are in the LT coding region (see 1 and 2 above the genome representation). The methods used to detect recombination are RDP (R) GENCONV (G), BOOTSCAN (B), MAXCHI (M), CHIMERA (C), SISCAN (S) and 3SEQ (T) with the methods with the lowest p-value being underlined.

## Concluding remarks

Eight polyomavirus genomes were recovered from three bark scorpions collected in urban backyards in Phoenix and Tempe, Arizona (USA). Four polyomavirus genomes were recovered from the livers and four from the guts of the sampled animals. All bark scorpion specimens appeared healthy at the time of collection and no investigation into possible associated disease states was undertaken. The eight polyomavirus genomes sampled from these species represent three distinct species. Genomes representing species 1 and 2 were isolated from two scorpions from both the liver and gut, whereas examples of species 3 were recovered from one scorpion, but from both the liver and gut. This is reminiscent of the situation for more carefully studied placental mammals, which typically harbor chronic co-infections with multiple polyomavirus species. While it is highly likely that the viruses recovered from the liver infect scorpions, it cannot be formally ruled out that the genomes recovered from the gut might reflect predation on other arachnids or insects. Even if scorpions are not the true host, since they prey only on other arthropods, these viruses are still likely to infect arthropods. Further investigation into the viromes of scorpions and other arachnids is likely to yield evidence of a broad range of novel polyomaviruses. Based on the recovery and characterization of these genomes, it appears that at least one polyomavirus was circulating among the last common ancestors of arthropods and chordates.

## Data availability

The sequences have been deposited in GenBank with accession #s MW175892-MW175899.

## Acknowledgements

The molecular work described here was supported with startup funds from Arizona State University made available to AV. This work was funded in part by the NIH Intramural Research Program, with support from the NCI Center for Cancer Research awarded to CB.

## Notes

### Competing Interest Statement

The authors have declared no competing interest.

